# An imputed ancestral reference genome for the *Mycobacterium tuberculosis* complex better captures structural genomic diversity for reference-based alignment workflows

**DOI:** 10.1101/2023.09.07.556366

**Authors:** Luke B Harrison, Vivek Kapur, Marcel A Behr

## Abstract

Reference-based alignment of short-reads is a widely used technique in genomic analysis of the *Mycobacterium tuberculosis* complex (MTBC) and the choice of reference sequence impacts the interpretation of analyses. The most widely used reference genomes include the ATCC type strain (H37Rv) and the putative MTBC ancestral sequence of Comas *et al*. both of which are based on a lineage 4 sequence. As such, these referents do not capture the complete structural variation now known to be present in the MTBC. To better represent the base of the MTBC, we generated an imputed ancestral genomic sequence, termed MTBC_0_ from reference-free alignments of closed MTBC genomes. When used as a reference sequence in alignment workflows, MTBC_0_ mapped more short sequencing reads and called more SNPs relative to the Comas et al. sequence while exhibiting minimal impact on the overall phylogeny of MTBC. The results also show that MTBC_0_ provides greater fidelity in capturing genomic variation and allows for the inclusion of regions absent in H37Rv such as the TbD1 and RvD4496/RD7/RD713 regions in standard MTBC workflows without additional steps. The use of MTBC_0_ as an ancestral reference sequence into a standard workflows modestly improved read mapping, SNP calling and intuitively facilitates the study of structural variation and evolution in MTBC.

**Data Summary:** The MTBC_0_ sequence, is available in the online data supplement in FASTA format at https://github.com/lukebharrison/MTBC0. Included with the MTBC_0_ sequence in the data supplement are: the reference-free alignment of MTBC closed genomes in hierarchical alignment (HAL) format, control files for cactus, annotations for H37Rv and L8, a BED file of regions excluded from SNP calls lifted over onto MTBC_0_, as well as the scripts used to call SNPs and the phylogenetic trees generated in this article. All previously published sequence data is available at the NCBI nucleotide and SRA databases, accession number for sequences used in this manuscript are available in Supplementary Tables 1 and 2.

**Impact Statement:** This article describes an imputed ancestral genomic sequence (MTBC_0_) at the base of the MTBC for use as a reference sequence for *Mycobacterium tuberculosis* genomic workflows. Widely used reference sequences are limited to the structural diversity present in H37Rv, a lineage 4 isolate. MTBC_0_ obviates this limitation by incorporating the structural variation present at the base of the *Mycobacterium tuberculosis* complex (MTBC) by encompassing a wide sample of human and animal lineages including newly discovered lineages (L8, *M. orygis*). Use of MTBC_0_ enables the mapping of more reads and calling of more SNPs and allows for the investigation of structural variation not present in the current used reference sequences within this important group of animal and human pathogens.

## Introduction

Tuberculosis in humans and animals is caused by infection with closely related bacteria that comprise the *Mycobacterium tuberculosis* complex (MTBC). Over the past decade, studies of the phylogeny, evolution and molecular epidemiology of the MTBC have been conducted using next generation sequencing (NGS) workflows. The vast majority of NGS workflows rely on a reference-based alignment of short sequencing reads to assemble genomic sequences, call single nucleotide polymorphisms (SNPs) and investigate structural variation. Current workflows used for MTBC (e.g. MTB-seq; Kohl et al., 2018) have used either the genome of the reference strain H37Rv (Cole et al., 1998) or an estimated most-recent common ancestor of the MTBC developed by Comas et al. (2010).

**Table 1.** Summary of mapping results and SNP calls of 310 short-read MTBC genomes using the MTBC_0_, Comas et al., and H37Rv reference sequences.

| Reference genome | Length (bp) | % reads mapped, median (IQR) | % reads unmapped, median (IQR) | Filtered SNPs called, median (IQR) |
| --- | --- | --- | --- | --- |
| MTBC0 | 4435783 | 99.7% (0.15) | 0.3% (0.15) | 513 (74) |
| Comas et al. | 4411532 | 99.3% (0.38) | 0.7% (0.38) | 488.5 (73.5) |
| H37Rv | 4411532 | 99.3% (0.33) | 0.7% (0.40) | 673.5 (597.5) |

Although widely used, the choice of H37Rv is not ideal: this genome is within lineage 4 of the MTBC and thus variation called against this genome represents the sum of evolution up the tree from H37Rv to the most-recent common ancestor (MRCA) of the MTBC and back down the lineage to the sequence in question. Further, as this is a tip-to-tip comparison, the directionality of any evolutionary change is not immediately resolved. Further, regions of deletion present in H37Rv (referred to as Rv-Deletions [RvDs] – Brosch et al., 2002) prevent mapping of reads from those regions, even if present in the genome under investigation. This latter issue has led to the use of workarounds based on realignment of unmapped reads to an alternative referent (e.g. *Mycobacterium canettii*, for example in Brites et al., 2018).

Recognizing these issues, Comas et al. (2010) proposed the use of an estimate of the MRCA of the MTBC (defined by the closed genome sampling available, using lineages L1-L6). This addressed the problem of a tip-to-tip comparison, but this sequence does not incorporate newly available genomic data and lineages (L7-9, animal lineages). Furthermore, this estimated MRCA is based on the structural variation present in the H37Rv genome and is thus unsuitable for the direct alignment of RvDs.

An ideal reference genome for the MTBC would 1) contain the structural variation present at the root of the MTBC to maximize mapping of reads, 2) represent the ancestral state of genomic positions to polarize evolutionary events informed by the recent discovery of deeply branching MTBC lineages (e.g. L8, Ngabonziza et al., 2020). Here, a new estimate of the ancestral state of the genome at the root of the MTBC is estimated and its use as a reference genome is demonstrated with a common workflow: generation of a reference-based SNP alignment and phylogenetic tree. Its ability to better capture structural variation absent in H37Rv is then demonstrated at the TbD1 and RD7/RD713/RvD4496 regions (Brosch et al., 2002).

## Methods

### Estimation of the ancestral genome of the MTBC

To estimate the ancestral genome of MRCA of the MTBC, 30 closed genomes available on the NCBI GenBank database were downloaded (Supplementary Table 1). Genomes were adjusted for circularity manually and softmasked for highly repetitive regions using RepeatMasker v.4.1.5 (Smit, 2015). The ProgressiveCactus genome alignment tool was used to align genomes and estimate ancestral genomes (Armstrong et al., 2020). The cactus algorithm requires a set phylogenetic tree, and so one was generated using a SNP alignment generated using Parsnp, executed with default parameters (Treangen et al. 2014). A phylogenetic tree was estimated from this SNP alignment using RAxML v8.2.12, with the Lewis correction for ascertainment (Stamatakis, 2018) with the interrelationships of major lineages constrained to accepted relationships from recent large phylogenomic studies (e.g. Ngabonziza et al., 2020; Coscolla et al., 2021; Vågene et al., 2022). The position of lineage 8 was collapsed in a polytomy with the two well supported major clades in the MTBC (L5,6,9,A1-4) and (L1-4,7).

### SNP calling

A sample of 309 MTBC genomes consisting of short reads was selected from the genomes used by Chiner-Oms et al. (2022): all genomes from less common lineages were included, along with sub-sampling (10 random genomes per sub-lineage) of the major MTBC lineages with extensive sub-sampling (2,4). SRA accession numbers are provided in Supplementary Table 2. Genomes identified as by Chiner-Oms as drug resistant were excluded. Raw reads were filtered and Illumina tags clipped using trimmomatic with parameters: MINLNE:20 SLIDINGWINDOW 5:20 TRAILING:10 (Bolger et al., 2014). Then, Kraken2 (Wood et al., 2019) was used to select only reads mapping to *Mycobacterium* and duplicate reads were removed using picard v2.23.3. Reads were aligned to a reference genome (MTBC_0_, Comas et al., or H37Rv) with bwa mem (Li, 2009). GATK v4.2.2.0 was then used to call and filter SNPs and indels using GenotypeGVCF and VariantFiltration programs (filter parameters: QD<2.0, FS>60.0, MQRankSum<-12.5, Low40MQ, MQ<40.0, ReadPosRankSum<-8.0, DP<10).

Finally, filtered SNPs and indels were further filtered to remove SNPs within repetitive and PE/PPE regions (as classified by Goig et al., 2020), as is standard in pipelines (e.g. MTBseq, Kohl et al., 2018), by translating annotations using the cactus alignments in BED format from H37Rv using the halLiftover command (Armstrong et al., 2020).

### Phylogenetic Analysis

Filtered SNPs were then used to construct a multiple sequence alignment of variable positions using samtools v1.13 (Li et al., 2009), and a phylogenetic tree was estimated using RaxML v8.2.12, with the Lewis correction for ascertainment (Stamatakis, 2018). A rapid tree search was combined with 1000 rapid bootstrap pseudoreplicates under the GTRCAT model, followed by final optimization with the GTRGAMMA model. Phylogenies were plotted using the cophyloplot and comparePhylo tools in the phytools and APE R packages, respectively (Revell, 2012; Paradis and Schliep, 2019).

### Visualization of TbD1 and RD7/RD713/RvD4966

The location of the TbD1 region (NCBI accession AJ426486.1; Brosch et al., 2002) was identified in the MTBC_0_ reference sequence using blastn (Altschul, 1990). The region of deletion was visualized using the Integrative Genomics Viewer (IGV; Robinson et al., 2011) with MTBC_0_ as the reference genome, and aligned short reads from a selection of genomes in the 309 genome data set with high coverage (1 per lineage other than L8 where both available genomes are included, see Figure 2 for SRA accessions). Reference annotations for the closed genomes H37Rv and lineage 8 (NCBI CP048071.1) are included for context and were mapped using the cactus alignments and the halLiftover command (Armstrong et al., 2020). The RD7/RD713/RvD4966 regions, as defined by Brosch et al. [2002], Mostowy et al. [2004], and Liu et al. [2023] respectively, were identified based on H37Rv annotations and visualized as above in IGV.

## Results

### MTBC_0_

The imputed ancestral genome of the MRCA of the MTBC, MTBC_0_ is a total of 4.436Mb in length, capturing structural variation of an additional ∼24kb relative to H37Rv (4.412Mb). When used to align short reads from a sample of 309 MTBC genomes and 1 *M. canettii* genome, MTBC_0_ maps a higher proportion of filtered reads relative to H37Rv or the Comas et al. ancestor (summarized in Table 1, complete mapping statistics in Supplementary Table 3). Using a GATK-based SNP calling pipeline with filtering for SNP quality and filtering out SNPs falling in low complexity regions as well as IS elements and PE/PPE genes, the use of MTBC_0_ as a referent calls a median of ∼24 more SNPs relative to Comas et al. More SNPs are called with H37Rv as a referent.

Phylogenetic analysis of SNP alignments generated using an identical pipeline, but with varying reference sequence shows only small differences in topology (Figure 1, Supplementary Figure 1) within groups, but with similar overall between lineage relationships. The topology of the phylogenetic tree is congruent with other recent analyses (e.g. Ngabonziza et al., 2020; Coscolla et al., 2021; Vågene et al., 2022); lineage 8 is the most deeply branching lineage, and there is a deep division separating a) lineages 5,6,9 and the “animal-adapted” lineages from b) *Mycobacterium tuberculosis sensu stricto* (lineages 1,2,3,4,7). Major bifurcations in the tree were well supported in the bootstrap analysis for all trees.

**Figure 1.**
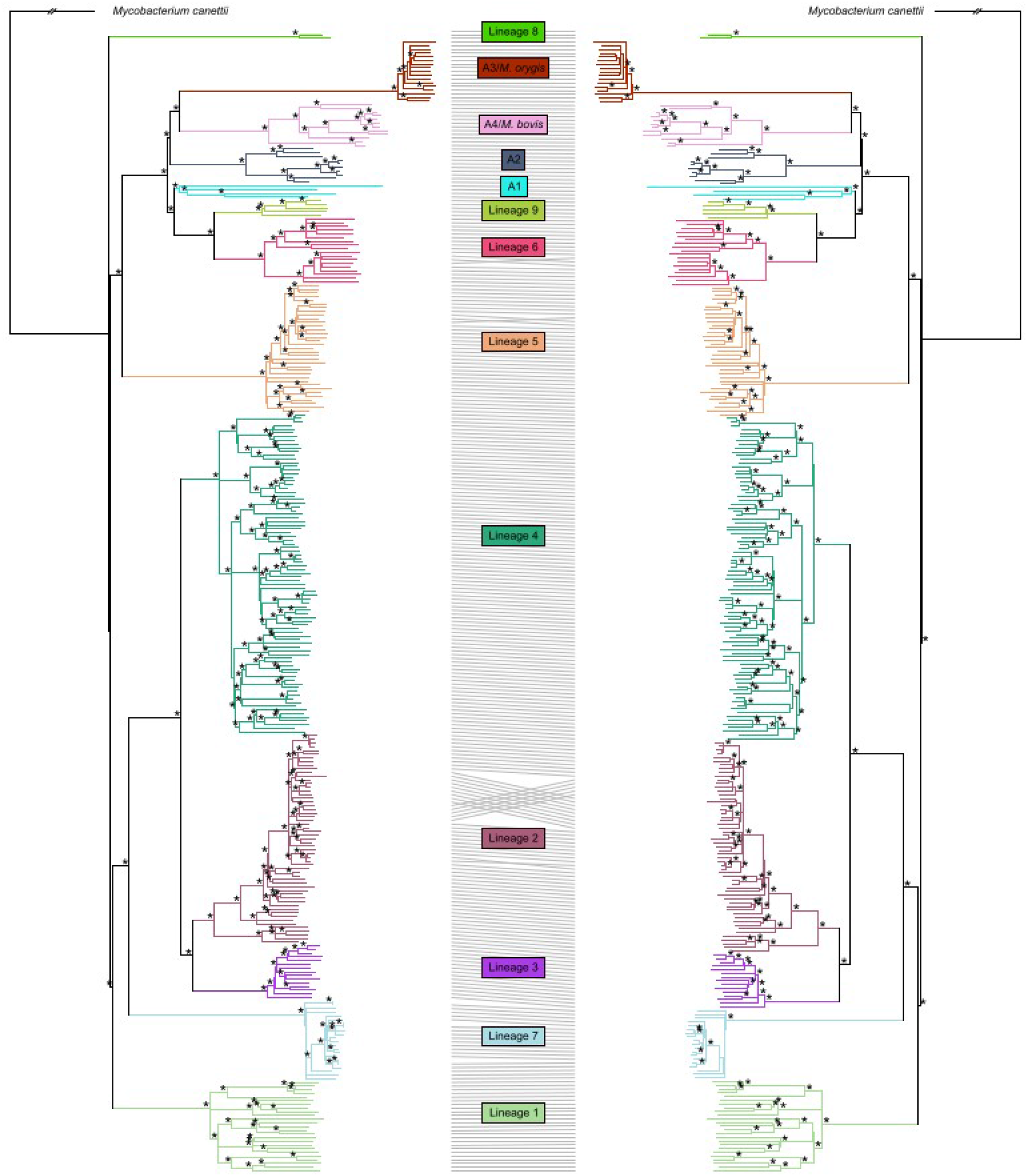
Maximum likelihood phylogeny trees of the MTBC based on SNPs called from short-read sequencing data by the GATK-based pipeline, using MTBC_0_ (left) and Comas et al., (2010) (right) reference sequences. Bootstrap values of 100% are indicated by *. Differences in phylogenetic tree topology are indicated by lines relating identical tips of the phylogeny.

The TbD1 region is visualized using against the MTBC_0_ sequence (Figure 2). An approximately 2.1Kb deletion is detected in the L2, 3 and 4 lineages. The RD7 region is similarly visualized (Figure 3) as expected in Lineage 6, 9 and A1-A4. The RD7 region also overlaps with RD713 in Lineage 5 and RvD4496 in Lineage 4; relative to MTBC_0_, these regions have lengths of: 4.4Kb, 5.9Kb and 17.2Kb, respectively.

**Figure 2.**
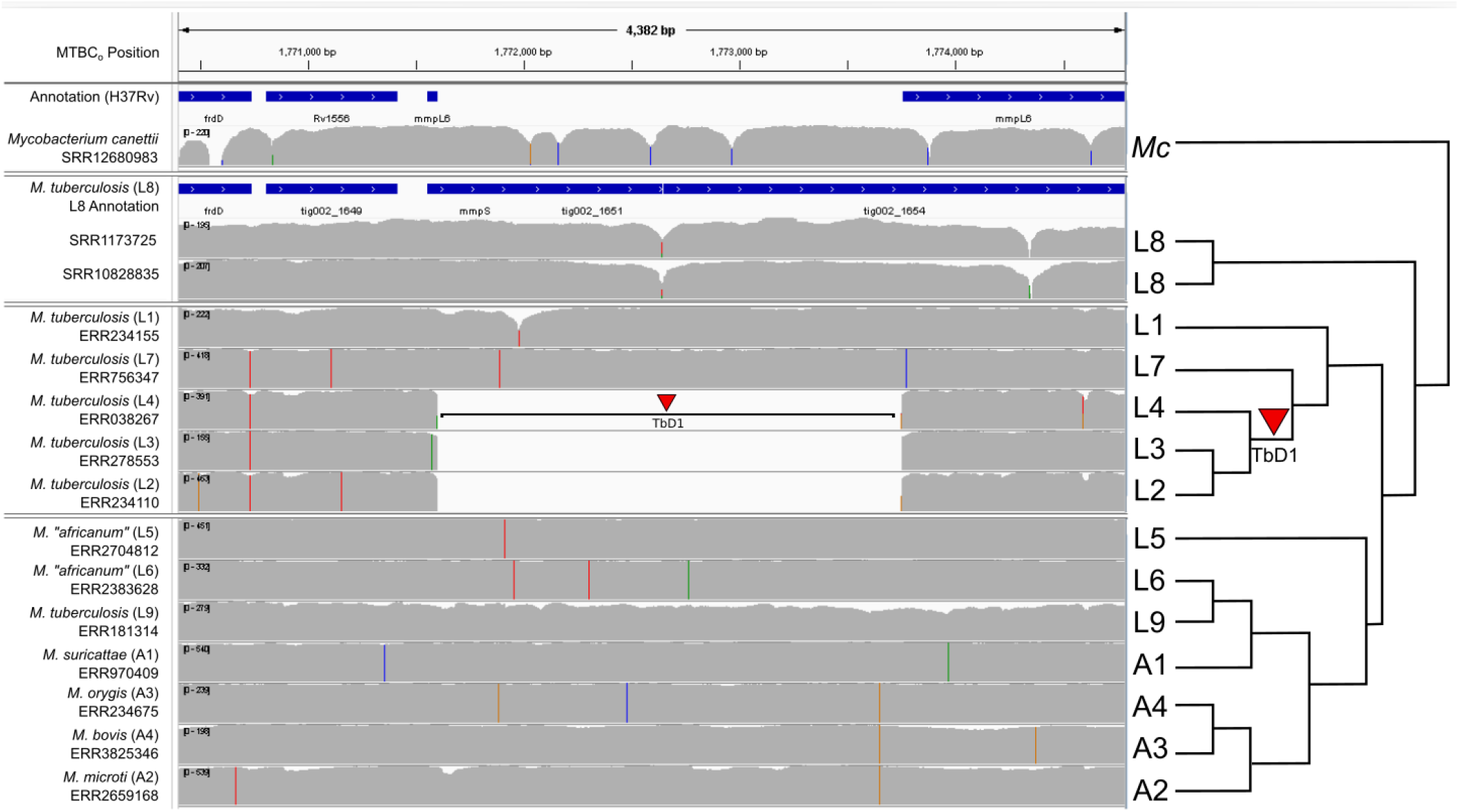
Summary of short-read alignments and log-transformed coverage for the TbD1 region relative to the MTBC_0_ reference sequence generated by IGV for a small sample of MTBC genomes (1 per lineage) and *M. canettii*. The figure is annotated on the right with a simplified phylogenetic tree topology of the MTBC estimated in Figure 1, with arbitrary branch lengths. Note the clear presence of a large ∼2.1Kb deletion in lineages 2, 3 and 4.

**Figure 3.**
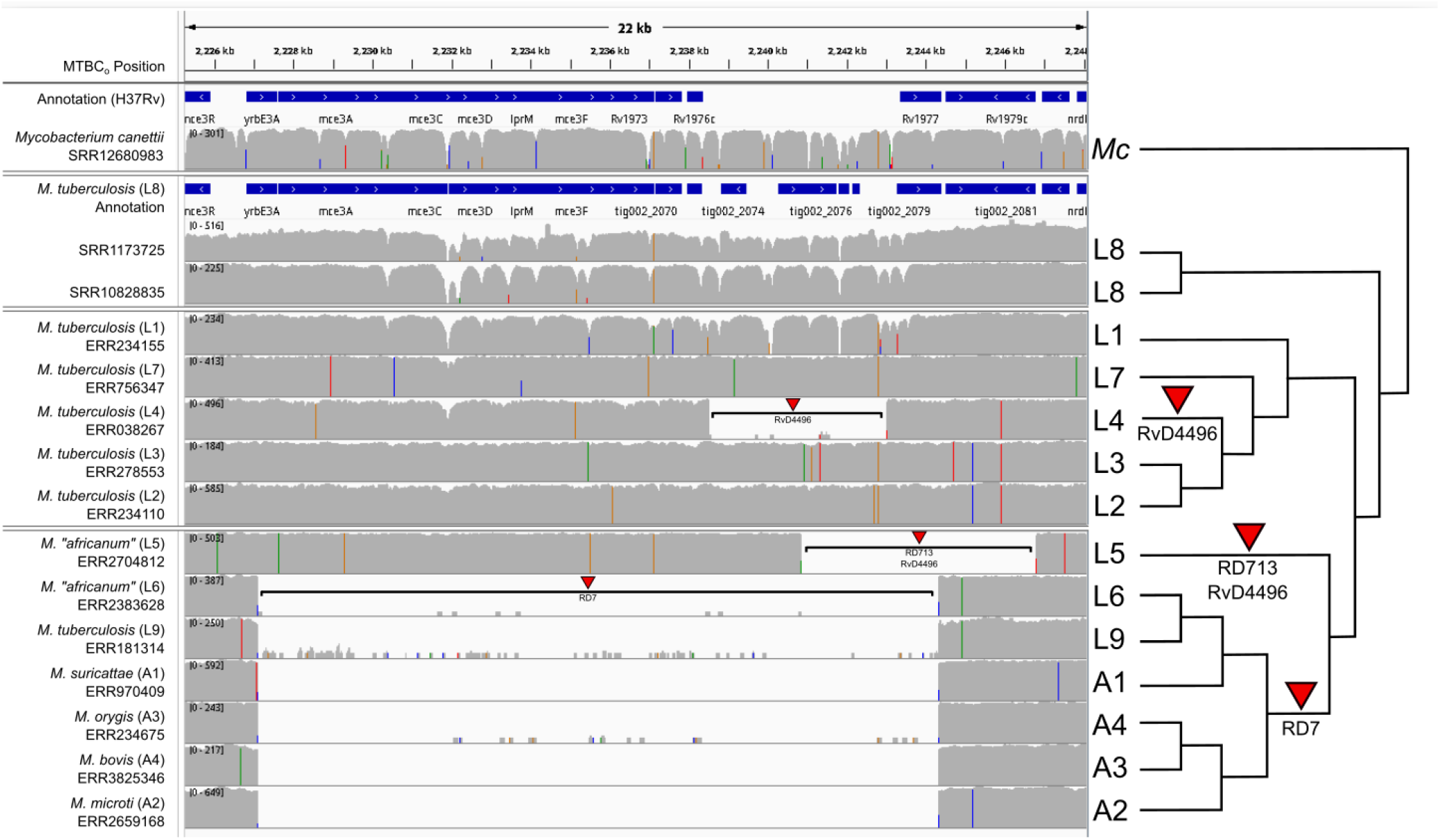
Summary of short-read alignment and log-transformed coverage for the region containing RD7, RD713, and RvD34496 relative to the MTBC_0_ reference sequence generated by IGV for a small sample of MTBC genomes (1 per lineage other than L8) and *M. canettii*. The figure is annotated on the right with a simplified phylogenetic tree topology of the MTBC estimated in Figure 1, with arbitrary branch lengths. The RD713 described by Mostowy et al. (2004) overlaps a deletion in H37Rv (RvD4496): using the H37Rv referent, RD713 is described as ∼3.7Kb, although a larger region is affected: ∼5.9Kb when MTBC_0_ is used as a referent. Likewise, the deleted region in RD7 is a 12.7Kb deletion relative to H37Rv, but a 17.2Kb deletion relative to MTBC_0_.

## Discussion

The use of MTBC_0_ as a reference sequence incrementally improves mapping of reads and SNP calling relative to Comas et al. sequence, albeit with only a subtle effect on a phylogeny estimated from these SNPs. As expected from a tip-to-tip comparison, using H37Rv results in more SNP calls as this represents the evolutionary distance from the MRCA of both genomes to both H37Rv and the genome of interest. However, the inferred phylogenetic tree is very similar (Supplementary Figure 1), and is also congruent with recent analyses (e.g. Ngabonziza et al., 2020). This is similar to previous work demonstrating only subtle effects on phylogeny of reference sequence choice within the MTBC (Lee and Behr, 2016). Although phylogenetically informative SNPs may exist in RvDs, additional information to further address fundamental questions in the evolutionary history of the MTBC, such as whether the MRCA was human or animal-adapted, which is still an open question, will likely come from further sampling and discovery of rare and deeply branching MTBC lineages.

Relative to the Comas et al., the use of MTBC_0_ captures structural variation that is not present in H37Rv and clearly identifies TbD1, an evolutionarily significant deletion specific to lineages 2, 3 and 4 (Bottai et al., 2020). It further demonstrates RvD4966, and clarifies the size of the overlapping RD7 and RD713 deletions, which are underestimated relative to H37Rv given the concomitant RvD.

Previous studies using the reference sequences of H37Rv or Comas et al., have relied on supplementary workflows examining unmapped reads followed by alignment against *M. canettii*. Although RvDs can be characterized in this manner, their position is then reported relative to one of the diverse *M. canettii* genomes (e.g. Liu et al, 2022). Other analyses may also benefit from the use of the MTBC_0_ as a reference sequence. These include the phylogenetic placement of genomes from ancient DNA analysis or from to be discovered deeply branching MTBC clades, and the intuitive analysis of regions of deletions present. A final potential application is fine-grained molecular epidemiology analysis of clades distant from H37Rv that may benefit from the marginally increased SNP resolution and incorporation of RvDs.

This approach has several drawbacks. The structural variation present in MTBC_0_ is based on the alignment of 30 closed genomes and is thus limited to variation present in that sample. The discovery of additional deeply branching lineages similar to what was found with L8 may reveal additional regions for future consideration. Further, as it is an estimate of the ancestral genome present at the base of the MTBC, the structural variation represented in MTBC_0_ is limited to that estimated to be present at the root of the complex. This latter concern is somewhat mitigated by the paucity of reported insertions (other than IS elements) or horizontal gene transfer events in the evolutionary history of the MTBC (Gagneux, 2018).

In the longer term, the continued development and refinement of long-read-based third generation sequence technologies may enable the widespread use of reference-free workflows (e.g. EnteroBase; Achtman et al., 2022) that rely on *de novo* assembly. However, until the availability of long-read based genomic data approaches that contained in the vast databases of short-read genomic sequences available and being generated for the MTBC, reference-based methods are likely to predominate. MTBC_0_ is designed to complement H37Rv and Comas *et al*.’s reference sequences as another tool in the toolbox, perhaps as a new ‘North star’ to facilitate genomic analyses in the MTBC, particularly for studies that need to capture the evolution of and within structural variation absent in H37Rv. Further, although not comprehensively examined here, MTBC_0_ provides a new estimate of the ancestral genomic states, both in terms of gene content and sequence. These permit intuitive interpretation of evolutionary changes and may inform estimates of ancestral phenotypic parameters in the search for the origin of the *Mycobacterium tuberculosis* complex.

## Ethical approval and consent to participate

n/a.

## Author contributions

LBH, MAB, and VK conceptualized the project, LBH performed the analyses and wrote the original draft of the article. LBH, MAB, and VK reviewed and revised the article. MAB provided resources for the project. MAB and VK supervised the project.

## Acknowledgments

The authors would like to thank Tod Stuber from the USDA-Veterinary Services for thoughtful comments and discussion on an early version the MTBC_0_ sequence.

## Supplementary Materials

**Supplementary Tables (in a single spreadsheet file, with three tables in individual tabs as below)**

**Supplementary Table 1**. Closed genomes used to impute MTBC_0_.

**Supplementary Table 2**. Genomic data used in this analysis.

**Supplementary Table 3**. Detailed results of short read alignments and SNP filtering pipeline for each reference sequence (MTBC_0_, Comas et al., H37Rv)

**Supplementary References**. Included as an additional tab in the supplementary tables file, with references for materials cited in Supplementary tables.

**Supplementary Figure 1.**
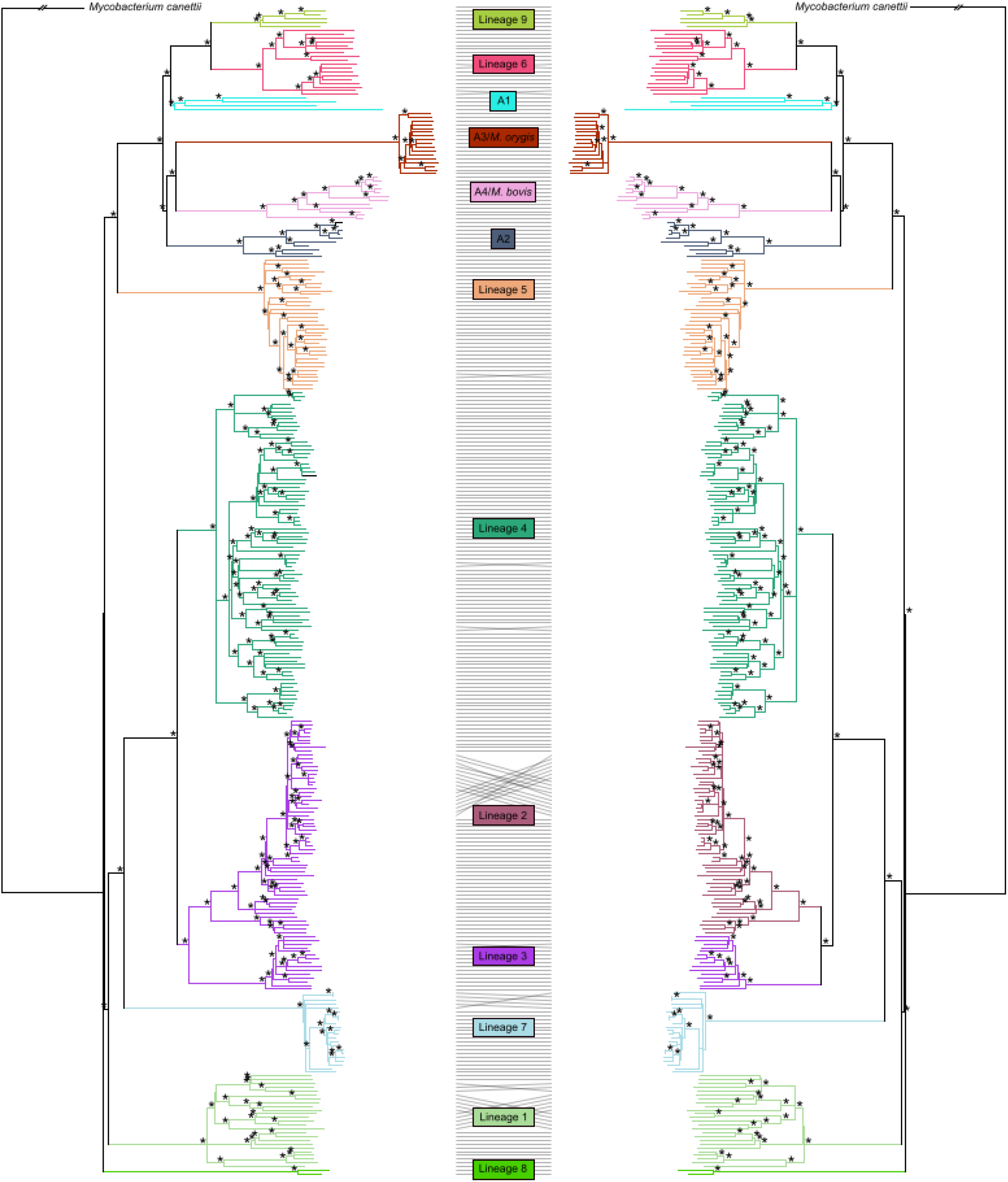
Maximum likelihood phylogeny trees of the MTBC based on SNPs called from short-read sequencing data by the GATK-based pipeline, using MTBC_0_ (left) and H37Rv (right) reference sequences. Bootstrap values of 100% are indicated by *. Differences in phylogenetic tree topology are indicated by lines relating identical tips of the phylogeny.

